# AnnoLnc: a web server for systematically annotating novel human lncRNAs

**DOI:** 10.1101/042655

**Authors:** Mei Hou, Xing Tang, Feng Tian, Fangyuan Shi, Fenglin Liu, Ge Gao

## Abstract

Although the repertoire of human lncRNAs has rapidly expanded, their biological function and regulation remain largely elusive. Here, we present AnnoLnc (http://annolnc.cbi.pku.edu.cn), an online portal for systematically annotating newly identified human lncRNAs. AnnoLnc offers a full spectrum of annotations covering genomic location, RNA secondary structure, expression, transcriptional regulation, miRNA interaction, protein interaction, genetic association and evolution, as well as an abstraction-based text summary and various intuitive figures to help biologists quickly grasp the essentials. In addition to an intuitive and mobile-friendly Web interactive design, AnnoLnc supports batch analysis and provides JSON-based Web Service APIs for programmatic analysis. To the best of our knowledge, AnnoLnc is the first web server to provide on-the-fly and systematic annotation for newly identified human lncRNAs. Some case studies have shown the power of AnnoLnc to inspire novel hypotheses.

## Introduction

Long noncoding RNAs (lncRNAs) are operationally defined as RNA transcripts that are 1) longer than 200 nt and 2) do not encode proteins (Rinn and Chang 2012). With high-throughput screening and follow-up experimental validation, several studies show that lncRNAs play essential roles in almost every important biological process, including imprinting (Lee and Bartolomei 2013), cell cycles (Kitagawa et al. 2013), tumorigenesis (Park et al. 2014) and pluripotency maintenance (Ng et al. 2012) through multiple mechanisms, such as guides, scaffolds, and decoys, as well as chromatin architecture organizers (Trimarchi et al. 2014; Jalali et al. 2015).

In recent years, the repertoire of human lncRNAs has rapidly expanded. Approximately 50% of human lncRNAs in the GENCODE catalog were identified in the past five years (15,512 in GENCODE v7 increased to 28,031 in GENCODE v24) (Derrien et al. 2012; Harrow et al. 2012). A recent study identified more than 30,000 additional unannotated human lncRNAs genes (Iyer et al. 2015). However, the functional roles of lncRNAs remain largely elusive: less than 1% of identified human lncRNAs have been experimentally investigated (Quek et al. 2015), driving the need for computational methods.

Several studies have proposed methods for *in silico* prediction of the function of novel lncRNAs. The “guilt-by-association” strategy is the most widely used approach (Guttman et al. 2009). A dedicated web server, ncFAN, was developed to predict lncRNA functions based on enriched functional terms of coding genes in the same co-expression module (Liao et al. 2011a; Liao et al. 2011b); the algorithm was improved by taking protein-protein interaction into account (Guo et al. 2013). Moreover, several attempts have been made to characterize molecules interacting with a given lncRNA (Agostini et al. 2013; Lu et al. 2013; Li et al. 2014; He et al. 2015; Suresh et al. 2015).

The immense diversity of lncRNAs’ functions calls for an integrative annotation pipeline that incorporates multiple disparate genome-scale datasets, offering a broad functional spectrum for novel human lncRNAs (Jalali et al. 2015). Here we present AnnoLnc, a one-stop portal for systematically annotating novel human lncRNAs (see **Figure 1**b for a detailed comparison with similar tools). Based on more than 700 data sources and various tool chains, AnnoLnc accepts human lncRNA sequences as input, enabling a systematic annotation encompassing genomic location, secondary structure, expression patterns, transcriptional regulation, miRNA interaction, protein interaction, genetic association and evolution. An intuitive web interface is available for interactive analysis, supporting both desktops and mobile devices. Users can upload multiple sequences and perform batch analysis in one click. The results will be further summarized automatically in plain English for quick reading. Programmers can further integrate AnnoLnc into their pipeline through standard JSON-based Web Service APIs.

**Figure 1.**
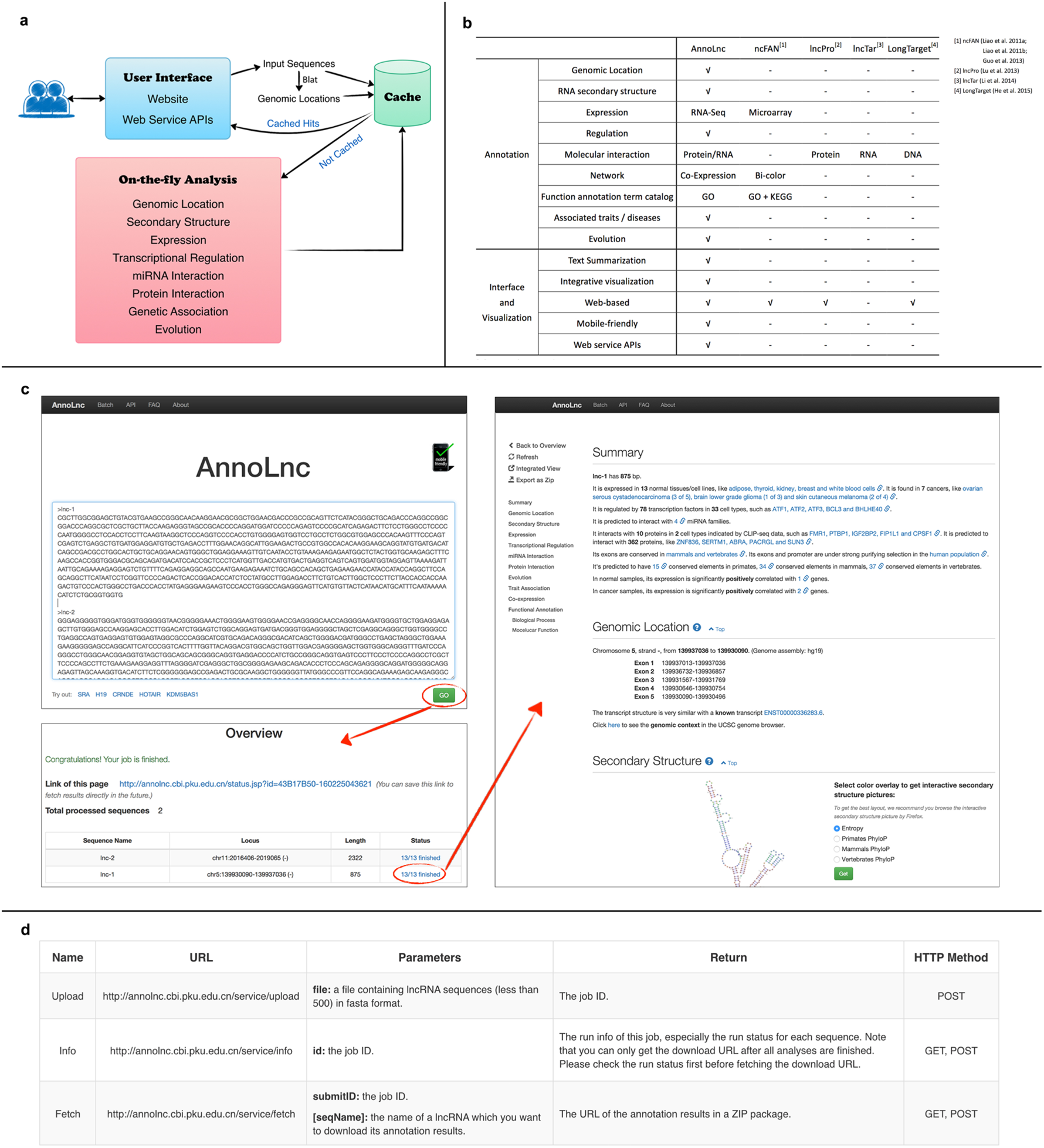
An overall introduction to AnnoLnc. **a)** The architecture of the AnnoLnc web server. Users can submit RNA sequences on the AnnoLnc website or by Web service APIs. First, AnnoLnc tries to map input sequences to the human genome and, if possible, obtain genomic locations. Then, AnnoLnc searches the global cache for possible hits based on both sequences and aligned splicing structures. Cached results are returned directly if a hit is detected, and novel sequences and loci are sent to on-the-fly analysis modules. **b)** Comparison of AnnoLnc with other similar tools. AnnoLnc has the most comprehensive annotations and is much better at the interface and visualization. **c)** AnnoLnc web pages. Only a few clicks are required from submission of sequences to receipt of annotation results. **d)** The introduction to Web service APIs provided by AnnoLnc.

## Results and Discussion

### Systematic annotation of human lncRNAs

Designed as a flexible platform, AnnoLnc provides systematic annotation of human lncRNAs through various modules and summarizes the results as figures, tables as well as plain English (**Figure 1**a and **1**c).

#### Genomic location

When a lncRNA is submitted online, AnnoLnc first identifies its genomic coordinate and splicing structure by aligning the input sequence to the human reference genome. The coordinates are further compared with annotated human gene models compiled from lncRNAdb (Quek et al. 2015) and GENCODE (Harrow et al. 2012) (see Materials and Methods for details), and a direct link to the corresponding database entry is provided for hits.

#### Secondary structure

The structure of an RNA molecule is essential for its biological functions. For each input lncRNA, the secondary structure is folded based on the minimum free energy principle (Lorenz et al. 2011) and then rendered online as an interactive plot.

Biological functions of secondary structures lead to local stability and bring evolutionary constraints onto the sequences of lncRNAs (Smith et al. 2013). T o h e l p u s e r s identify functional motifs, AnnoLnc allows users to color each base in the structure plot by its corresponding entropy or conservation score (**Figure 2**a and **2**b).

**Figure 2.**
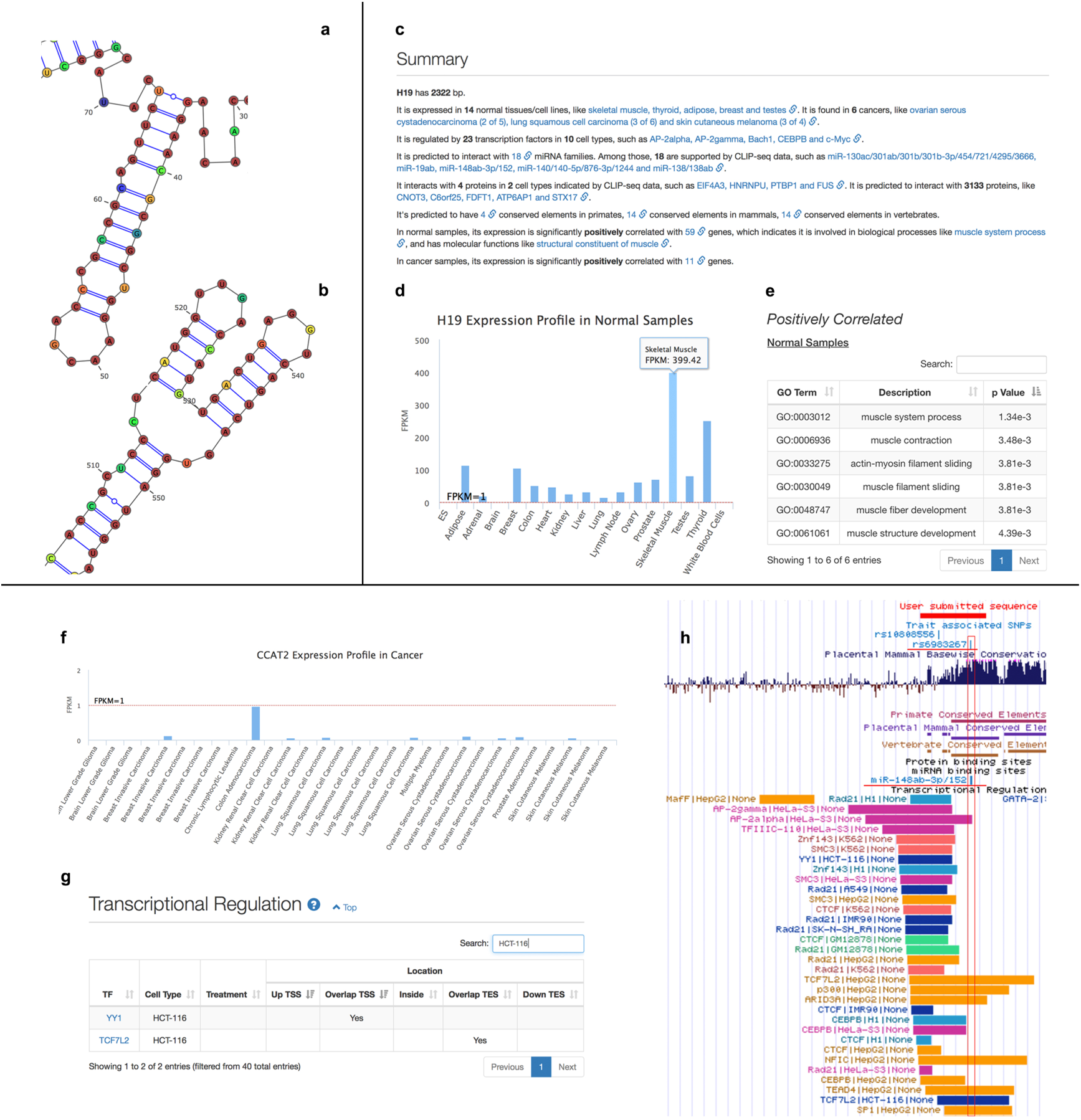
Case studies for AnnoLnc. **a-b)** The case study of lncRNA *SRA*. In the interactive secondary structure plot with vertebrate phyloP scores as the color overlay, two sub-structures are very easily to be identified because most bases are colored red. **c-e)** The case study of lncRNA *H19*. **c** is a summary of the annotation results, which helps users quickly grasp the essentials. **d** is the expression profile of *H19* in normal samples. It has the highest expression in “skeletal muscle”. **e** is the predicted GO terms based on positively correlated coding genes in normal samples. The terms are all muscle related. **f-h)** The case study of lncRNA *CCAT2*. **f** is the expression profile of *CCAT2* in cancer cell lines. It has the highest expression in “colon adenocarci-noma”. **g** is the result of “transcriptional regulation”. Searching the CRC cell line “HCT-116” shows that *CCAT2* is regulated by TCF7L2 and YY1. Ling *et al.* has verified that *CCAT2* and TCF7L2 can regulate each other and form a feedback loop to promote CRC (Ling et al. 2013). **h** is the integrative view in the UCSC genome browser for annotations at the transcript level. It is easy to determine that rs6983267 is within the seed binding site of the miRNA family miR-148ab-3p/152.

#### Expression profile

A transcript’s expression pattern also provides important hints about its functionality (Guttman et al. 2009). For each input lncRNA, the expression profiles in 17 normal samples (16 adult healthy tissues and one embryonic stem cell line) and 30 cancer cell lines (covering 10 common cancers) are presented in interactive charts (**Figure 2**d and **Figure 2**f). T o improve the response time, the expression of known lncRNAs (including lncRNAs in GENCODE v19 and lncRNAdb v2) was pre-calculated and loaded into the global cache. For novel lncRNAs, we adopt the LocExpress method to perform on-the-fly expression estimation accurately and efficiently (see Materials and Methods for more details).

To help users identify co-regulated partners of the input lncRNA, AnnoLnc reports co-expressed genes based on normal samples and cancer samples. An expression-based functional prediction is further performed by identifying statistically significant enriched Gene Ontology (GO) terms based on co-expressed protein-coding genes. Adjusted *p* values for the multiple-testing issue are reported as well (**Figure 2**e) (Guttman et al. 2009; Tang et al. 2013).

#### Transcriptional regulation

Transcriptional factors largely determine the expression level of lncRNAs. Integrating 498 ChIP-Seq datasets, AnnoLnc locates the binding sites of 159 TFs in the input lncRNA locus, and reports these binding sites based on their relative location to the lncRNA locus, such as “upstream transcriptional start site (TSS)”, “overlap with TSS”, “inside the lncRNA loci”, “overlap with transcriptional end site (TES)” and “downstream TES” (**Table S1**a, **Figure 2**g).

#### miRNA interaction

Interacting with miRNAs, lncRNAs can be post-transcriptionally regulated or act as decoys (Yoon et al. 2014). AnnoLnc provides predicted miRNA family partners of lncRNAs and highlights high-confidence binding sites based on conservation scores of multiple clades and 61 integrated CLIP-Seq datasets (for details about CLIP-Seq dataset processing, refer to the “Protein interaction” section). If a predicted site is supported by CLIP-Seq data, it is marked “CLIP support” (**Table S1**b).

#### Protein interaction

lncRNAs can interact with multiple proteins, as guides and/or scaffolds, to perform their functions (Wang and Chang 2011). CLIP-Seq is one of the most widely used high-throughput methods to detect RNA-protein interactions experimentally (McHugh et al. 2014). AnnoLnc screens putative protein partners for an input lncRNA in 112 CLIP-Seq datasets covering 51 RNA binding proteins. In case of methodology bias introduced by heterogeneous analysis pipelines, all the CLIP-Seq data were reanalyzed locally with a uniform pipeline (see Materials and Methods for more details). Finally, protein partners, cell types, treatments and corresponding *p* values reported by the analysis pipeline are reported to users.

In addition, AnnoLnc conducts *in silico* prediction across the entire human proteome for each lncRNA by lncPro (Lu et al. 2013). To improve the specificity, we estimate *p* value for each hit based on an empirical NULL distribution. Then, the predicted protein partners, interaction scores and empirical *p* values are reported.

#### Genetic association

Large-scale genetic association studies enable detection of multiple phenotypic traits that lncRNAs may associate with (Cheetham et al. 2013). By integrating the NHGRI GWAS Catalog (Welter et al. 2014), AnnoLnc links an input lncRNA to diseases/traits based on strong linkage blocks defined by linkage disequilibrium (LD) values in multiple populations. Then, linked SNPs, corresponding tagSNPs, traits/diseases, *p* values, significance, LD values and populations from which these LD values are derived, as well as supporting PubMed IDs, are reported to users (**Table S**2).

#### Evolution

The evolutionary signature is an important hint as to biological function. AnnoLnc generates an interactive overview chart for users to easily interrogate inter-species conservation of exons and promoter regions among primate, mammal and vertebrate clades. To further detect intra-species purifying selection, AnnoLnc also provides the derived allele frequency for input lncRNAs based on 1000 Genomes Project data (Genomes Project et al. 2010). Because many lncRNAs are partially conserved, we also report conserved elements predicted by phastCons (Siepel et al. 2005) in different clades with the length and score, which is an indicator of conservation. These conserved blocks can help users identify functional elements combined with other annotation results, especially in the integrated plot.

### User interface and visualization

AnnoLnc is designed to be intuitive. The most common operations (such as submitting sequences and obtaining annotation results) can be performed with just a few clicks (**Figure 1**c). The default Web interface is implemented based on responsive design, which enables the optimal view for both desktop PCs and mobile devices. T o further help users quickly grasp the essentials from abundant annotations generated by various modules, AnnoLnc provides a concise summary text in plain English for each input lncRNA at the top of the annotation result page by abstraction-based summarization, with inline links available for checking original results when necessary (**Figure 2**c). Furthermore, to help users explore the genomic context of input lncRNAs, AnnoLnc supports exporting transcript-level annotations (including transcript structure, TF binding sites, miRNA binding sites, protein binding sites and SNPs) onto the UCSC genome browser as pre-tuned custom tracks (**Figure 2**h).

In addition to the browser-based interactive analysis, AnnoLnc provides a “batch mode” that allows users to upload multiple sequences together and fetch annotations as a ZIP. AnnoLnc also offers a set of JSON-based web service APIs to help advanced users run the analysis and fetch results programmatically, enabling an easy integration of AnnoLnc into downstream analysis pipelines (see http://annolnc.cbi.pku.edu.cn/api.jsp for more detailed instruction as well as the demo code).

### Case studies by AnnoLnc

The noncoding form of the steroid receptor RNA activator (*SRA*, AF092038, http://annolnc.cbi.pku.edu.cn/cases/SRA) has been reported to function as a noncoding RNA by Lanz *et al.* (Lanz et al. 1999) and is the first lncRNA that has experimentally derived secondary structure, which was derived by Novikova *et al.* (Novikova et al. 2012). In the interactive secondary structure plot with vertebrate phyloP score as color overlay in AnnoLnc, it is easy to identify two conserved regions. One is a hairpin region from base 30 to 72 (**Figure 2**a). With approximately 75% of bases colored red, this conserved sub-structure is clearly distinguishable from others. In fact, this region corresponds to the most conserved H2 sub-structure highlighted by Novikova *et al.* Site-directed mutagenesis of this region reduced the co-activation performance of *SRA* by 40% (Lanz et al. 2002), suggesting the importance of lncRNA secondary structure on its function (Mercer and Mattick 2013). The other distinct region is a three-way junction hairpin sub-structure from base 506 to 555 with 78% colored red (**Figure 2**b). This region is very similar to the conserved regions H15, H16 and H17 verified by Novikova *et al.*

*H19* (http://annolnc.cbi.pku.edu.cn/cases/H19) is the first identified imprinting lncRNA (Brannan et al. 1990; Bartolomei et al. 1991). Consistent with the work by Dey *et al.* (Dey et al. 2014), AnnoLnc shows that *H19* has the highest expression in “skeletal muscle” (**Figure 2**d) and is associated with muscle-related function terms such as “muscle fiber development” (GO:0048747, **Figure 2**e). Moreover, the “transcriptional regulation” module reports that *H19* is regulated by multiple known cell proliferation-and cell cycle-related TFs, including c-Myc, Max, Maz, and E2F6 in cancer cell lines (**Table S1**a), confirming its previously reported tumorigenesis function (Guo et al. 2014). In addition, AnnoLnc identified 18 CLIP-Seq-supported binding miRNA families (**Table S1**b), and several miRNAs have already been verified experimentally, such as miR-138 in colorectal cancer (Liang et al. 2015) and miR-17-5p in HeLa cells and myoblasts (Imig et al. 2015).

In addition to confirming previous reports, the integrative annotations provided by AnnoLnc help users to generate new hypotheses. For example, lncRNA *CCAT2* (http://annolnc.cbi.pku.edu.cn/cases/CCAT2) has been reported to promote colorectal cancer (CRC) growth and metastasis, and risk allele G of rs6983267 within the *CCAT2* transcript is associated with up-regulated expression of the lncRNA (Ling et al. 2013). Integrating miRNA annotation with the variant track (**Figure 2**h), SNP rs6983267 is found to be just within the seed binding site of miRNA miR-148ab-3p/152, suggesting that SNP rs6983267 might weaken the binding of miR-148ab-3p/152 and increases the transcript level of *CCAT2*.

## Conclusion

T o the best of our knowledge, AnnoLnc is the first online web server to systematically annotate novel human lncRNAs. Compared with similar tools (**Figure 1**b), the annotation generated by AnnoLnc covers a much wider range of perspectives with intuitive visualization and summarization. Several case studies have shown the power of AnnoLnc to systematically annotate lncRNAs and inspire novel hypotheses for follow-up experimental studies. Employing Web Service APIs, AnnoLnc is friendly for not only interactive users, but also programmers for batch analysis. In the future, we plan to upgrade the AnnoLnc server by adding more analysis modules, including RNA-DNA interactions as well as literature mining, and further improving the integration with more intuitive and interactive facilities.

## Materials and Methods

### Annotation analysis

As a Web server, AnnoLnc runs on-the-fly analysis for every input sequence.

#### Mapping input sequences to the reference genome

All input sequences are aligned against the reference human genome hg19 with Blat (Kent 2002). When a single sequence is aligned in multiple places, genome-wide best alignments are identified by standard pslSort and pslReps. In case of false-positive junction sites caused by mismatches or small indels, putative exons shorter than 20 bp (as well as putative introns shorter than 40 bp) are discarded.

#### Secondary structure

RNAfold v2.0.7 in the ViennaRNA package (Lorenz et al. 2011) is employed to predict secondary structures, with the option “--noLP” enabled to avoid undesirable isolated base pairs. When multiple candidates are available, the one with minimum free energy is kept, as recommended by the authors of the ViennaRNA package.

#### Expression-and co-expression-based functional annotation

The expression profile of input transcripts is estimated based on 64 RNA-Seq datasets (covering normal adult tissues, tumor cell lines as well as human embryonic stem cell lines; see **Table S3** for more details). We mapped the reads of 34 normal tissue/cell line samples to human genome hg19 by TopHat (v1.4.1.1). Bam files of 30 cancer samples were downloaded directly from CGHub. (see **Table S3** and **S4** for the number of mapped reads and CGHub IDs, respectively.)

##### Pre-calculated expression profiles of known transcripts

We generated a gene model (GM) gtf file (http://annolnc.cbi.pku.edu.cn/about/annolnc_gene_model_v1.gtf.gz) covering human lncRNAs in lncRNAdb v2.0 (Quek et al. 2015) based on GENCODE (Harrow et al. 2012) v19. First, we downloaded human lncRNA sequences in the lncRNAdb and obtained transcript structures as described under “Mapping input sequences to the reference genome”.

These transcript structures were compared with GENCODE v19 by Cuffcompare (v2.1.1). If the code was “=” or “c”, the lncRNA was replaced by the known transcript; otherwise it was considered a “novel transcript” and merged into GENCODE v19. The expression of all annotated transcripts in the GM file was pre-calculated by StringTie (v1.0.4) with the options “-e-b”, and then normalized by the geometric method in normal and cancer samples separately.

##### On-the-fly expression estimation of input transcripts

Taking advantage of the local nature of RNA-Seq, we developed a novel quantification method called LocExpress for real-time estimation of the expression level based on pre-mapped reads (Hou *et al*., manuscript in preparation). Briefly, LocExpress infers the minimum spanning bundle (MSB) of an input transcript based on its genomic coordinate as well as reads mapped in the corresponding region. Then, the estimated relative abundance is further adjusted and normalized based on the original size (*i.e*., the total mass derived from all mapped reads) and reported in canonical FPKM unit. In our preliminary assessment, the LocExpress method usually takes less than one minute for one novel lncRNA across all samples, and the result is highly consistent with that of the classical method (http://annolnc.cbi.pku.edu.cn/images/LocExpress.png). For the normal sample set, the FPKM of two replicates of each tissue/cell line are averaged to report to users.

##### Co-expression analysis

Thirty-four normal samples and thirty cancer samples were treated separately. To avoid the duplicated GO annotation for isoforms, we first obtained expression profiles at the gene level by adding the FPKM of all transcripts of each gene in the GM file. Then, we filtered these genes as described below, resulting in 29,798 genes in the normal sample set and 25,449 genes in the cancer sample set.

1. FPKM filter. The sum of FPKM in all samples should be not less than 1.
2. Tissue-specific filter. The tissue-specific score is calculated by the “getsgene” function in the R package rsgcc. If a gene has a score larger than 0.85, it is not considered fit for the co-expression analysis.

For submitted transcripts that pass the above filters, the Pearson correlations with genes are calculated. Then, highly correlated genes are reported by a “gradually decreased” criterion to remove putative false positives and retain true positives. If there are more than 10 genes with r ≥ 0.9, GO enrichment analysis is performed with these genes directly. If not, we determine whether there are 10 genes above the cutoff of 0.8. This process continues until the cutoff arrives at 0.7. Negatively correlated genes are identified in a similar manner. GO enrichment analysis for these correlated genes is further conducted with the R package GOstats, and significantly enriched GO terms (adjusted *p* value ≤ 0.01) are reported as putative functional assignments of the input transcript.

#### Transcriptional regulation

AnnoLnc integrated 498 ChIP-Seq datasets covering 159 transcript factors (TFs) in 45 cell xslines (see **Table S5** for more details). Uniform peak files generated by the ENCODE project were downloaded from http://hgdownload.cse.ucsc.edu/goldenPath/hg19/encodeDCC/wgEncodeAwgTfbsUniform/. For each input transcript, AnnoLnc searches putative TF binding sites within 5 Kb upstream and 1 Kb downstream and reports all sites based on their position relative to the transcript.

#### miRNA interaction

TargetScan v6.0 (Friedman et al. 2009) is employed to search putative miRNA binding sites. T o reduce the potential false positive rate, we run the prediction on 87 highly conserved miRNA families (**Table S6**) derived from miRcode (Jeggari et al. 2012) (http://www.mircode.org/download.php). Then, conservation scores in primate, mammal and vertebrate clades for each identified site are calculated as described in Jeggari *et al.* For example, 10 species in primates are used in the Target Scan prediction, and if a binding site is identified in eight species, the conservation score in primates is 8/10 = 0.8. In mammals and vertebrates, scores are calculated in the same manner except that “mammals” are “non-primate mammals” (26 species) and “vertebrates” are “non-mammal vertebrates” (10 species). To further highlight high-confidence sites, predicted sites are screened based on a pre-compiled 61 AGO CLIP-Seq dataset (**Table S7**, see the “ Calling RNA-protein interactions based on CLIP-Seq data

” section for more details about CLIP-Seq analysis), and hit sites are considered “CLIP supported”.

#### Protein interaction

##### Calling RNA-protein interactions based on CLIP-Seq data

A total of 112 CLIP-Seq datasets were integrated in AnnoLnc, covering 51 RNA binding proteins (RBPs) other than AGO (see **Table S8** for a full list) from the GEO. All these data were reanalyzed uniformly in case of methodology bias. Briefly, we first trimmed the adapter by FASTX Clipper, and only reads longer than 15 nt were kept and mapped to human genome hg19 by the algorithm BWA-backtrack (v0.7.10-r789) (Li and Durbin 2009) with the options “-n 1-i 0” (allow one alignment error). Then, only unique mapped reads were kept. To improve precision, we used stringent criteria for site calling with PIPE-CLIP v1.0.0 (Chen et al. 2014); FDR cutoffs for both enriched clusters and reliable mutations were set as 0.05 (cross-linking sites in HITS-CLIP data identified by deletion, insertion and substitution were combined).

To evaluate the performance of our pipeline, we downloaded raw reads of wild-type FET proteins (FUS, EWSR1 and TAF15) from DDBJ (SRA025082) and performed the analysis described above. For comparison with reported results (Supplementary Data 1, (Hoell et al. 2011)), cross-linking sites identified by both methods were mapped to RefSeq IDs. Our pipeline shows fairly high precision (0.95 for FUS, 0.96 for EWSR1, and 0.91 for TAF). Meanwhile, we evaluated on a HITS-CLIP dataset for the DGCR8 protein (Macias et al. 2012). We downloaded raw reads of all four samples (D8.1, D8.2, T7.1 and T7.2) from GEO (GSE39086) and analyzed them as described above (D8.1 data was excluded because PIPE-CLIP failed to generate cross-linking sites with a “model failed to converge” error). Comparison with the original results downloaded from http://regulatorygenomics.upf.edu/Data/DGCR8/ also shows good precision (0.89 for D8.2, 0.74 for T7.1, and 0.78 for T7.2).

##### Ab initio prediction of lncRNA-protein interaction

lncPro (Lu et al. 2013) is employed for *ab initio* prediction of lncRNA-protein interactions in the whole human proteome. We downloaded 99,459 human protein sequences from Ensembl, filtered 1,917 sequences that could not be processed by lncPro (containing “*”, “X”, “U” or length not within 30-30,000 AA), ultimately obtaining 97,542 protein sequences. For efficiency, we modified the source code of lncPro to pre-calculate all protein features in batch. To improve specificity, we further derived the statistical significance of the interaction scores reported by lncPro based on empirical NULL distribution (http://annolnc.cbi.pku.edu.cn/images/lncPro.png) generated by random shuffling. Interactions with a *p* value ≤ 0.01 are considered to be significant. To make the results more intuitive, Ensembl IDs are finally converted into HGNC gene symbols. If multiple Ensembl IDs are mapped to one gene symbol, the score with the smallest *p* value are reported.

#### Genetic association

To To exhaustively detect genetic association, AnnoLnc first scans all SNPs within the transcript region (5 Kb upstream to 1 Kb downstream of each input transcript). Using one of these SNPs as an example, a SNP is linked with a tagSNP if it is within the haplotype region (defined as r^2^ > 0.5, ftp://ftp.ncbi.nlm.nih.gov/hapmap/ld_data/2009-04_rel27/) tagged by the tagSNP reported in the NHGRI GWAS Catalog (Welter et al. 2014) (downloaded from the UCSC genome browser). Then, this linked SNP, corresponding tagSNPs, traits/diseases, p values, significance (defined as *p* value ≤ 5e-8), LD values, populations from which these LD values are derived, as well as supporting PubMed IDs, are reported by AnnoLnc.

#### Evolution

For each submitted lncRNA, we incorporated the 46 way phyloP score (primates, mammals/placentals and vertebrates) from the UCSC Genome Browser, and the derived allele frequency (DAF) (Genomes Project et al. 2010) of the YRI population (Yoruba in Ibadan, Nigeria) from http://compbio.mit.edu/human-constraint/ for every position (if has corresponding scores) of both the exon and promoter region (1 Kb upstream). To obtain an overall view, we also calculate the mean scores for the exon and promoter regions and organized the scores into bar charts. The phastCons conserved elements were downloaded from ftp://hgdownload.cse.ucsc.edu/goldenPath//hg19/database. The score reported to users is the LOD score. Conserved elements shorter than 20 bp are omitted from the table.

### AnnoLnc website

The AnnoLnc website runs on the Tomcat server. The backend is based on Java Servlet and MySQL database. In the frontend, some JavaScript libraries are used to facilitate accessibility. Bootstrap is used for the mobile friendly layout. jQuery is used for Ajax. DataTables is used to show tables and Highcharts for interactive charts. The display of the interactive SVG plot is enabled by “svg-pan-zoom,” available at https://github.com/ariutta/svg-pan-zoom.

## Acknowledgements

*Funding*: This work was supported by funds from the China 863 Program (2015AA020108), the Seeding Grant for Medicine and Life Sciences of Peking University (2014-MB-13), and the State Key Laboratory of Protein and Plant Gene Research. The research of G.G. was supported in part by the National Program for Support of Top-notch Young Professionals. Part of the analysis was performed on the Computing Platform of the Center for Life Sciences of Peking University.

